## Supplementary Material for "Rate of brain aging associates with future executive function in Asian children and older adults"

### Supplementary Methods

#### Finetuning Process and Details

Due to the high computational cost of model training, we employed an heuristic hyperparameter search. We started with GUSTO as it had the worst initial performance. We consulted the original model's author (Esten Leonardsen), who suggested tuning the last layer alone with dropout = 0.3, weight decay = 1e-3, and learning rate = 1e-3. We originally stopped model training once the validation MAE failed to improve after 3 epochs in a row. However, when tuning only the last layer, we found that the performance did not meaningfully improve and the learning curves showed evidence of underfitting (Supplementary Figure S6A).

Thus, we next tried to tune all layers with the same hyperparameters. This resulted in much better performance, but the learning curves showed some evidence of instability (Supplementary Figure S6B). Since Leonardsen previously observed instability with high learning rates, we tried to anneal the learning rate using cosine decay. Based on the shape of the previous training curve (Supplementary Figure S6B), we set the number of decay steps to 25 epochs. This resulted in better performance and more stability (Supplementary Figure S6C).

Having gotten satisfactory performance on GUSTO, we then tried applying the same hyperparameters to EDIS and SLABS. However, in these datasets, we observed that the model changed too quickly, leading to a very high validation error at the beginning of training and ultimately worse performance (Supplementary Figure S6D). This was likely because the original pretrained model did not need to adjust as much to fit these datasets. Thus, we tried to reduce the initial learning rate by factors of 10, observing better training and performance at 1e-4 and 1e-5 (Supplementary Figure S6E). We observed underfitting and worse performance at 1e-6 (Supplementary Figure S6F). Thus, we optimized the initial learning rate out of {1e-3, 1e-4, 1e-5} for each fold in each dataset. Observing that the model had well-converged by 35 epochs, we stopped training at 35 epochs. Since this resulted in good performance, we kept all other hyperparameters constant.

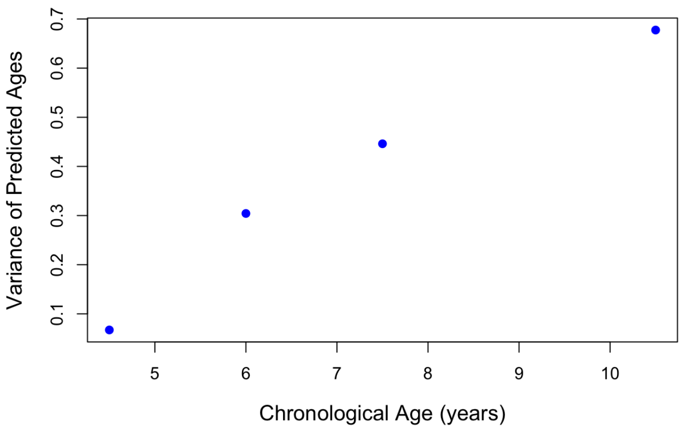


Figure S1: Variance of **finetuned** predicted ages by age group in GUSTO

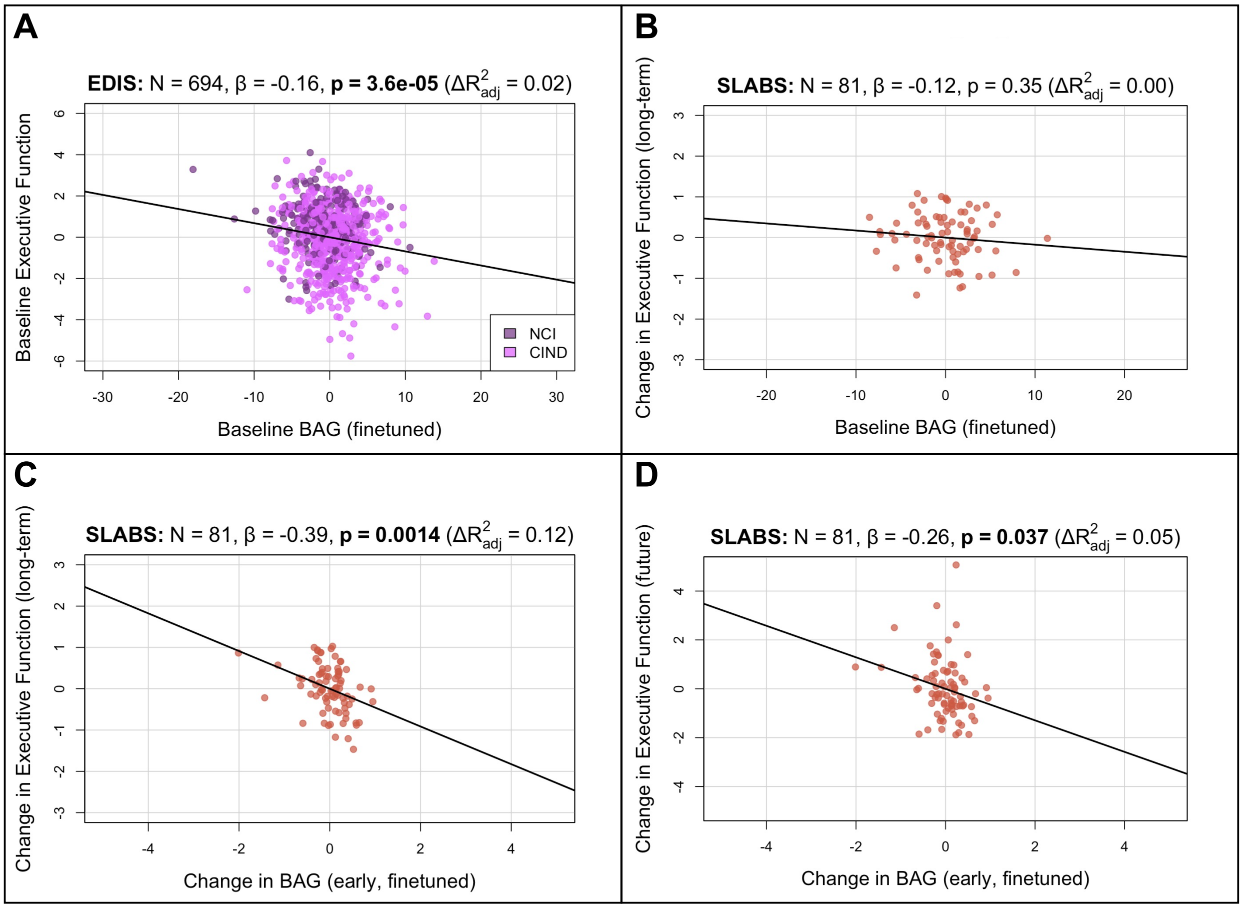


Figure S2: Brain age gap from the **finetuned** model remains negatively associated with executive function in **elderly**. Compare to Figure 3 of the main text.

| Characteristic | EDIS (N = 694) |
|---|---|
| Global cognition z-score | $-2.55 \pm 2.21$ ($-10.47$ – 1.96) |
| Executive function domain z-score | $-1.57 \pm 1.85$ ($-7.79$ – 1.26) |
| Attention domain z-score | $-2.13 \pm 2.63$ ($-13.69$ – 2.65) |
| Language domain z-score | $-1.84 \pm 1.70$ ($-11.72$ – 2.85) |
| Visuomotor speed domain z-score | $-1.64 \pm 1.76$ ($-5.87$ – 2.25) |
| Visuoconstruction domain z-score | $-2.50 \pm 2.57$ ($-13.63$ – 2.37) |
| Verbal memory domain z-score | $-1.40 \pm 1.40$ ($-5.17$ – 2.78) |
| Visual memory domain z-score | $-1.50 \pm 1.48$ ($-9.10$ – 2.15) |
| **Characteristic** | **SLABS (N = 81)** |
| Global cognition T-score | $51.82 \pm 4.75$ (41.21 – 61.65) |
| Executive function domain T-score | $51.98 \pm 5.57$ (42.39 – 64.08) |
| Attention domain T-score | $50.52 \pm 6.11$ (39.62 – 65.04) |
| Processing speed domain T-score | $52.96 \pm 7.22$ (36.13 – 71.84) |
| Verbal memory domain T-score | $52.18 \pm 7.70$ (32.38 – 67.04) |
| Visuospatial memory domain T-score | $51.48 \pm 7.62$ (35.00 – 65.80) |
| Cognition follow up (years) | $7.83 \pm 0.97$ (5.58 – 9.59) |
| **Characteristic** | **GUSTO (N = 217 to 239)** |
| KBIT-2 Composite IQ Standard Score (N = 217, 4.5 years old) | $92.38 \pm 14.24$ (52 – 132) |
| WCST Total Errors Standard Score (N = 220, 8.5 years old) | $99.45 \pm 15.95$ (64 – 136) |
| NEPSY-II Naming Scaled Score (N = 239, 8.5 years old) | $10.45 \pm 3.66$ (3 – 19) |
| NEPSY-II Inhibition Scaled Score (N = 239, 8.5 years old) | $10.31 \pm 3.3$ (2 – 18) |
| NEPSY-II Switching Scaled Score (N = 239, 8.5 years old) | $9.26 \pm 4.03$ (1 – 19) |

Table S1: Participant cognitive characteristics at baseline. EDIS was cross-sectional, while SLABS and GUSTO were longitudinal. **Key:** EDIS – Epidemiology of Dementia in Singapore; SLABS – Singapore Longitudinal Aging Brain Study; GUSTO - Growing Up in Singapore Towards healthy Outcomes; KBIT-2 - Kaufman Brief Intelligence Test Second Edition; WCST - Wisconsin Card Sorting Test; NEPSY-II: A Developmental Neuropsychological Assessment Second Edition

| Fold | EDIS | SLABS | GUSTO |
|---|---|---|---|
| 0 | 1e-4 | 1e-4 | 1e-3 |
| 1 | 1e-4 | 1e-4 | 1e-3 |
| 2 | 1e-4 | 1e-5 | 1e-3 |
| 3 | 1e-4 | 1e-5 | 1e-3 |
| 4 | 1e-5 | 1e-4 | 1e-3 |
| 5 | 1e-4 | 1e-4 | 1e-3 |
| 6 | 1e-5 | 1e-5 | 1e-3 |
| 7 | 1e-4 | 1e-5 | 1e-3 |
| 8 | 1e-4 | 1e-5 | 1e-3 |
| 9 | 1e-4 | 1e-4 | 1e-3 |

Table S2: Initial learning rates for each dataset and fold. **Key:** EDIS – Epidemiology of Dementia in Singapore; SLABS – Singapore Longitudinal Aging Brain Study; GUSTO – Growing Up in Singapore Towards healthy Outcomes

| Age | Dataset | Label | Equation |
|---|---|---|---|
| Elderly | EDIS | I. Baseline vs. baseline | $\text{Cog}_{bl} = \text{BAG}_{bl} + \text{age} + \text{sex} + \text{education}$ |
| Elderly | SLABS | II. Baseline vs. change | $\Delta\text{Cog}_{long} = \text{BAG}_{bl} + \text{age} + \text{sex} + \text{education}$ |
| Elderly | SLABS | III. Change vs. change (overlapping) | $\Delta\text{Cog}_{long} = \Delta\text{BAG}_{early} + \text{BAG}_{bl} + \text{age} + \text{sex} + \text{education}$ |
| Elderly | SLABS | IV. Change vs. change (future) | $\Delta\text{Cog}_{future} = \Delta\text{BAG}_{early} + \text{BAG}_{bl} + \text{age} + \text{sex} + \text{education}$ |
| Children | GUSTO | I. Baseline vs. baseline | $\text{Cog}_{4.5} = \text{BAG}_{4.5} + \text{age} + \text{sex}$ |
| Children | GUSTO | II. Baseline vs. future | $\text{Cog}_{8.5} = \text{BAG}_{4.5} + \text{age} + \text{sex}$ |
| Children | GUSTO | III. Change vs. future | $\text{Cog}_{8.5} = \Delta\text{BAG}_{4.5-7.5} + \text{BAG}_{4.5} + \text{age} + \text{sex}$ |

Table S3: Model equations for analyzing associations with cognition. **Key:** Cog - standardized cognitive score; BAG – brain age gap; bl - baseline; Δ - annual rate of change

| **Variable of Interest** | **Outcome** | $\beta$ $(\mathbf{CI_L}, \mathbf{CI_U})$ | **p** | $\mathbf{p_{corr}}$ | $\mathbf{\Delta R^2_{adj}}$ | $\mathbf{R^2}$ |
|---|---|---|---|---|---|---|
| Baseline BAG (pretrained) | Baseline Global Cognition | -0.1125 (-0.17, -0.06) | $<$ **0.0001** | **0.0006** | 0.0106 | 0.5011 |
| Baseline BAG (pretrained) | Baseline Executive Function | -0.1029 (-0.17, -0.04) | **0.0019** | **0.0076** | 0.0085 | 0.3297 |
| Baseline BAG (pretrained) | Baseline Attention | -0.0404 (-0.11, 0.03) | 0.2461 | 0.2461 | 0.0004 | 0.2562 |
| Baseline BAG (pretrained) | Baseline Language | -0.1145 (-0.18, -0.05) | **0.0009** | **0.0047** | 0.0107 | 0.2677 |
| Baseline BAG (pretrained) | Baseline Visuomotor Speed | -0.0825 (-0.14, -0.02) | **0.0052** | **0.0136** | 0.0053 | 0.4684 |
| Baseline BAG (pretrained) | Baseline Visuo-construction | -0.0896 (-0.15, -0.03) | **0.0045** | **0.0136** | 0.0063 | 0.3911 |
| Baseline BAG (pretrained) | Baseline Verbal Memory | -0.1096 (-0.17, -0.05) | **0.0006** | **0.0034** | 0.0099 | 0.3826 |
| Baseline BAG (pretrained) | Baseline Visual Memory | -0.1395 (-0.20, -0.08) | $<$ **0.0001** | **0.0002** | 0.0165 | 0.3515 |

Table S4: Pretrained baseline vs. baseline results from EDIS. p-values are bolded if less than $\alpha = 0.05$. **Key:** BAG – brain age gap; $\beta$ – standardized regression coefficient; $CI_L$ - lower limit of 95% confidence interval; $CI_U$ - upper limit of 95% confidence interval; $p$ – uncorrected p-value; $p_{corr}$ – corrected p-value; $\Delta R^2_{adj}$ - change in adjusted $R^2$ when adding variable of interest; $R^2$ - model coefficient of determination

| Variable of Interest | Outcome | $\beta$ ($\mathbf{CI_L}, \mathbf{CI_U}$) | p | $\mathbf{p_{corr}}$ | $\mathbf{\Delta R^2_{adj}}$ | $\mathbf{R^2}$ |
|---|---|---|---|---|---|---|
| Baseline BAG (finetuned) | Baseline Global Cognition | -0.1661 (-0.23, -0.10) | $<$ **0.0001** | $<$ **0.0001** | 0.0171 | 0.5076 |
| Baseline BAG (finetuned) | Baseline Executive Function | -0.1607 (-0.24, -0.08) | $<$ **0.0001** | **0.0002** | 0.0158 | 0.3369 |
| Baseline BAG (finetuned) | Baseline Attention | -0.0632 (-0.14, 0.02) | 0.1223 | 0.1223 | 0.0015 | 0.2573 |
| Baseline BAG (finetuned) | Baseline Language | -0.1535 (-0.23, -0.07) | **0.0002** | **0.0007** | 0.0142 | 0.2712 |
| Baseline BAG (finetuned) | Baseline Visuomotor Speed | -0.1071 (-0.17, -0.04) | **0.0020** | **0.0040** | 0.0066 | 0.4697 |
| Baseline BAG (finetuned) | Baseline Visuocon-struction | -0.1291 (-0.20, -0.06) | **0.0005** | **0.0015** | 0.0099 | 0.3946 |
| Baseline BAG (finetuned) | Baseline Verbal Memory | -0.1722 (-0.24, -0.10) | $<$ **0.0001** | $<$ **0.0001** | 0.0183 | 0.3910 |
| Baseline BAG (finetuned) | Baseline Visual Memory | -0.2194 (-0.29, -0.15) | $<$ **0.0001** | $<$ **0.0001** | 0.0302 | 0.3651 |

Table S5: Finetuned baseline vs. baseline results from EDIS. p-values are bolded if less than $\alpha = 0.05$. **Key:** BAG – brain age gap; $\beta$ – standardized regression coefficient; $CI_L$ - lower limit of 95% confidence interval; $CI_U$ - upper limit of 95% confidence interval; $p$ – uncorrected p-value; $p_{corr}$ – corrected p-value; $\Delta R^2_{adj}$ - change in adjusted $R^2$ when adding variable of interest; $R^2$ - model coefficient of determination

| **Variable of Interest** | **Outcome** | $\beta$ $(\mathbf{CI_L}, \mathbf{CI_U})$ | **p** | $\mathbf{p_{corr}}$ | $\mathbf{\Delta R^2_{adj}}$ | $\mathbf{R^2}$ |
|---|---|---|---|---|---|---|
| Baseline BAG (pretrained) | Baseline Global Cognition | -0.0574 (-0.18, 0.06) | 0.3376 | 1.0000 | -0.0002 | 0.3514 |
| Baseline BAG (pretrained) | Baseline Executive Function | -0.0447 (-0.16, 0.07) | 0.4600 | 1.0000 | -0.0015 | 0.3379 |
| Baseline BAG (pretrained) | Baseline Verbal Memory | -0.0089 (-0.14, 0.12) | 0.8925 | 1.0000 | -0.0038 | 0.2090 |
| Baseline BAG (pretrained) | Baseline Visual Memory | -0.1299 (-0.27, 0.01) | 0.0716 | 0.4297 | 0.0105 | 0.0643 |
| Baseline BAG (pretrained) | Baseline Attention | -0.0187 (-0.15, 0.11) | 0.7750 | 1.0000 | -0.0035 | 0.2261 |
| Baseline BAG (pretrained) | Baseline Processing Speed | -0.0036 (-0.12, 0.11) | 0.9492 | 1.0000 | -0.0029 | 0.4038 |

Table S6: Pretrained baseline vs. baseline results from SLABS. p-values are bolded if less than $\alpha = 0.05$. **Key:** BAG – brain age gap; $\beta$ – standardized regression coefficient; $CI_L$ - lower limit of 95% confidence interval; $CI_U$ - upper limit of 95% confidence interval; $p$ – uncorrected p-value; $p_{corr}$ – corrected p-value; $\Delta R^2_{adj}$ - change in adjusted $R^2$ when adding variable of interest; $R^2$ - model coefficient of determination

| **Variable of Interest** | **Outcome** | $\beta$ $(\mathbf{CI_L}, \mathbf{CI_U})$ | **p** | $\mathbf{p_{corr}}$ | $\mathbf{\Delta R^2_{adj}}$ | $\mathbf{R^2}$ |
|---|---|---|---|---|---|---|
| Baseline BAG (finetuned) | Baseline Global Cognition | -0.0674 (-0.19, 0.05) | 0.2784 | 1.0000 | 0.0006 | 0.3522 |
| Baseline BAG (finetuned) | Baseline Executive Function | -0.0378 (-0.16, 0.09) | 0.5477 | 1.0000 | -0.0021 | 0.3373 |
| Baseline BAG (finetuned) | Baseline Verbal Memory | -0.0548 (-0.19, 0.08) | 0.4245 | 1.0000 | -0.0014 | 0.2113 |
| Baseline BAG (finetuned) | Baseline Visual Memory | -0.1243 (-0.27, 0.02) | 0.0974 | 0.5846 | 0.0081 | 0.0621 |
| Baseline BAG (finetuned) | Baseline Attention | 0.0056 (-0.13, 0.14) | 0.9342 | 1.0000 | -0.0038 | 0.2258 |
| Baseline BAG (finetuned) | Baseline Processing Speed | -0.0229 (-0.14, 0.09) | 0.7008 | 1.0000 | -0.0025 | 0.4042 |

Table S7: Finetuned baseline vs. baseline results from SLABS. p-values are bolded if less than $\alpha = 0.05$. **Key:** BAG – brain age gap; $\beta$ – standardized regression coefficient; $CI_L$ - lower limit of 95% confidence interval; $CI_U$ - upper limit of 95% confidence interval; $p$ – uncorrected p-value; $p_{corr}$ – corrected p-value; $\Delta R^2_{adj}$ - change in adjusted $R^2$ when adding variable of interest; $R^2$ - model coefficient of determination

| **Variable of Interest** | **Outcome** | $\beta$ $(\mathbf{CI_L}, \mathbf{CI_U})$ | **p** | $\mathbf{p_{corr}}$ | $\mathbf{\Delta R^2_{adj}}$ | $\mathbf{R^2}$ |
|---|---|---|---|---|---|---|
| Baseline BAG (pretrained) | Change in Global Cognition | -0.1213 (-0.36, 0.12) | 0.3232 | 1.0000 | -0.0002 | 0.0211 |
| Baseline BAG (pretrained) | Change in Executive Function | -0.2477 (-0.48, -0.01) | **0.0406** | 0.2433 | 0.0424 | 0.0711 |
| Baseline BAG (pretrained) | Change in Verbal Memory | -0.1263 (-0.37, 0.11) | 0.2970 | 1.0000 | 0.0013 | 0.0490 |
| Baseline BAG (pretrained) | Change in Visual Memory | 0.0337 (-0.21, 0.28) | 0.7815 | 1.0000 | -0.0122 | 0.0338 |
| Baseline BAG (pretrained) | Change in Attention | 0.0253 (-0.21, 0.27) | 0.8345 | 1.0000 | -0.0125 | 0.0457 |
| Baseline BAG (pretrained) | Change in Processing Speed | -0.1032 (-0.35, 0.14) | 0.4015 | 1.0000 | -0.0039 | 0.0157 |

Table S8: Pretrained baseline vs. change results from SLABS. p-values are bolded if less than $\alpha = 0.05$. **Key:** BAG – brain age gap; $\beta$ – standardized regression coefficient; $CI_L$ - lower limit of 95% confidence interval; $CI_U$ - upper limit of 95% confidence interval; $p$ – uncorrected p-value; $p_{corr}$ – corrected p-value; $\Delta R^2_{adj}$ - change in adjusted $R^2$ when adding variable of interest; $R^2$ - model coefficient of determination

| **Variable of Interest** | **Outcome** | $\beta$ $(\mathbf{CI_L}, \mathbf{CI_U})$ | **p** | $\mathbf{p_{corr}}$ | $\mathbf{\Delta R^2_{adj}}$ | $\mathbf{R^2}$ |
|---|---|---|---|---|---|---|
| Baseline BAG (finetuned) | Change in Global Cognition | -0.0480 (-0.30, 0.20) | 0.7041 | 1.0000 | -0.0116 | 0.0103 |
| Baseline BAG (finetuned) | Change in Executive Function | -0.1165 (-0.36, 0.13) | 0.3531 | 1.0000 | -0.0017 | 0.0292 |
| Baseline BAG (finetuned) | Change in Verbal Memory | -0.1431 (-0.39, 0.10) | 0.2491 | 1.0000 | 0.0045 | 0.0520 |
| Baseline BAG (finetuned) | Change in Visual Memory | 0.0409 (-0.21, 0.29) | 0.7431 | 1.0000 | -0.0118 | 0.0342 |
| Baseline BAG (finetuned) | Change in Attention | 0.1232 (-0.12, 0.37) | 0.3193 | 1.0000 | 0.0001 | 0.0576 |
| Baseline BAG (finetuned) | Change in Processing Speed | -0.0125 (-0.26, 0.24) | 0.9214 | 1.0000 | -0.0134 | 0.0066 |

Table S9: Finetuned baseline vs. change results from SLABS. p-values are bolded if less than $\alpha = 0.05$. **Key:** BAG – brain age gap; $\beta$ – standardized regression coefficient; $CI_L$ - lower limit of 95% confidence interval; $CI_U$ - upper limit of 95% confidence interval; $p$ – uncorrected p-value; $p_{corr}$ – corrected p-value; $\Delta R^2_{adj}$ - change in adjusted $R^2$ when adding variable of interest; $R^2$ - model coefficient of determination

| **Variable of Interest** | **Outcome** | $\beta$ $(\mathbf{CI_L}, \mathbf{CI_U})$ | **p** | $\mathbf{p_{corr}}$ | $\mathbf{\Delta R^2_{adj}}$ | $\mathbf{R^2}$ |
|---|---|---|---|---|---|---|
| Change in BAG (pretrained) | Change in Global Cognition | -0.1415 (-0.39, 0.11) | 0.2689 | 1.0000 | 0.0033 | 0.0370 |
| Change in BAG (pretrained) | Change in Executive Function | -0.3807 (-0.61, -0.15) | **0.0017** | **0.0100** | 0.1100 | 0.1864 |
| Change in BAG (pretrained) | Change in Verbal Memory | -0.0312 (-0.28, 0.22) | 0.8056 | 1.0000 | -0.0125 | 0.0498 |
| Change in BAG (pretrained) | Change in Visual Memory | 0.0917 (-0.16, 0.34) | 0.4718 | 1.0000 | -0.0064 | 0.0405 |
| Change in BAG (pretrained) | Change in Attention | -0.1872 (-0.44, 0.06) | 0.1371 | 0.6857 | 0.0164 | 0.0736 |
| Change in BAG (pretrained) | Change in Processing Speed | -0.0868 (-0.34, 0.17) | 0.5001 | 1.0000 | -0.0074 | 0.0217 |

Table S10: Pretrained change vs. change results from SLABS. p-values are bolded if less than $\alpha = 0.05$. **Key:** BAG – brain age gap; $\beta$ – standardized regression coefficient; $CI_L$ - lower limit of 95% confidence interval; $CI_U$ - upper limit of 95% confidence interval; $p$ – uncorrected p-value; $p_{corr}$ – corrected p-value; $\Delta R^2_{adj}$ - change in adjusted $R^2$ when adding variable of interest; $R^2$ - model coefficient of determination

| **Variable of Interest** | **Outcome** | $\beta$ $(\mathbf{CI_L}, \mathbf{CI_U})$ | **p** | $\mathbf{p_{corr}}$ | $\mathbf{\Delta R^2_{adj}}$ | $\mathbf{R^2}$ |
|---|---|---|---|---|---|---|
| Change in BAG (finetuned) | Change in Global Cognition | -0.2367 (-0.48, 0.01) | 0.0576 | 0.2305 | 0.0360 | 0.0570 |
| Change in BAG (finetuned) | Change in Executive Function | -0.3861 (-0.62, -0.15) | **0.0014** | **0.0084** | 0.1190 | 0.1536 |
| Change in BAG (finetuned) | Change in Verbal Memory | -0.1017 (-0.35, 0.14) | 0.4091 | 1.0000 | -0.0041 | 0.0607 |
| Change in BAG (finetuned) | Change in Visual Memory | -0.0284 (-0.28, 0.22) | 0.8200 | 1.0000 | -0.0128 | 0.0349 |
| Change in BAG (finetuned) | Change in Attention | -0.2651 (-0.50, -0.03) | **0.0287** | 0.1434 | 0.0493 | 0.1163 |
| Change in BAG (finetuned) | Change in Processing Speed | -0.0842 (-0.33, 0.17) | 0.5050 | 1.0000 | -0.0076 | 0.0125 |

Table S11: Finetuned change vs. change results from SLABS. p-values are bolded if less than $\alpha = 0.05$. **Key:** BAG – brain age gap; $\beta$ – standardized regression coefficient; $CI_L$ - lower limit of 95% confidence interval; $CI_U$ - upper limit of 95% confidence interval; $p$ – uncorrected p-value; $p_{corr}$ – corrected p-value; $\Delta R^2_{adj}$ - change in adjusted $R^2$ when adding variable of interest; $R^2$ - model coefficient of determination

| **Variable of Interest** | **Outcome** | $\beta$ $(\mathbf{CI_L}, \mathbf{CI_U})$ | **p** | $\mathbf{p_{corr}}$ | $\mathbf{\Delta R^2_{adj}}$ | $\mathbf{R^2}$ |
|---|---|---|---|---|---|---|
| Baseline BAG (pretrained) | Future WCST Standard Score | -0.0585 (-0.24, 0.13) | 0.5334 | 0.9798 | -0.0028 | 0.0256 |
| Baseline BAG (pretrained) | Future Naming (NEPSY-II) | 0.1290 (-0.05, 0.31) | 0.1573 | 0.6292 | 0.0043 | 0.0149 |
| Baseline BAG (pretrained) | Future Inhibition (NEPSY-II) | 0.1074 (-0.07, 0.29) | 0.2387 | 0.7161 | 0.0017 | 0.0155 |
| Baseline BAG (pretrained) | Future Switching (NEPSY-II) | 0.0630 (-0.12, 0.24) | 0.4899 | 0.9798 | -0.0022 | 0.0106 |

Table S12: Pretrained baseline vs. future results from GUSTO. p-values are bolded if less than $\alpha = 0.05$. **Key:** BAG – brain age gap; $\beta$ – standardized regression coefficient; $CI_L$ - lower limit of 95% confidence interval; $CI_U$ - upper limit of 95% confidence interval; $p$ – uncorrected p-value; $p_{corr}$ – corrected p-value; $\Delta R^2_{adj}$ - change in adjusted $R^2$ when adding variable of interest; $R^2$ - model coefficient of determination

| **Variable of Interest** | **Outcome** | $\beta$ $(\mathbf{CI_L}, \mathbf{CI_U})$ | **p** | $\mathbf{p_{corr}}$ | $\mathbf{\Delta R^2_{adj}}$ | $\mathbf{R^2}$ |
|---|---|---|---|---|---|---|
| Baseline BAG (finetuned) | Future WCST Standard Score | -0.0046 (-0.20, 0.20) | 0.9642 | 1.0000 | -0.0046 | 0.0239 |
| Baseline BAG (finetuned) | Future Naming (NEPSY-II) | -0.0108 (-0.21, 0.19) | 0.9143 | 1.0000 | -0.0042 | 0.0065 |
| Baseline BAG (finetuned) | Future Inhibition (NEPSY-II) | 0.0829 (-0.11, 0.28) | 0.4086 | 1.0000 | -0.0013 | 0.0125 |
| Baseline BAG (finetuned) | Future Switching (NEPSY-II) | -0.0359 (-0.23, 0.16) | 0.7203 | 1.0000 | -0.0037 | 0.0091 |

Table S13: Finetuned baseline vs. future results from GUSTO. p-values are bolded if less than $\alpha = 0.05$. **Key:** BAG – brain age gap; $\beta$ – standardized regression coefficient; $CI_L$ - lower limit of 95% confidence interval; $CI_U$ - upper limit of 95% confidence interval; $p$ – uncorrected p-value; $p_{corr}$ – corrected p-value; $\Delta R^2_{adj}$ - change in adjusted $R^2$ when adding variable of interest; $R^2$ - model coefficient of determination

| **Variable of Interest** | **Outcome** | $\beta$ $(\mathbf{CI_L}, \mathbf{CI_U})$ | **p** | $\mathbf{p_{corr}}$ | $\mathbf{\Delta R^2_{adj}}$ | $\mathbf{R^2}$ |
|---|---|---|---|---|---|---|
| Change in BAG (pretrained) | Future WCST Standard Score | -0.0639 (-0.21, 0.08) | 0.3877 | 1.0000 | -0.0011 | 0.0290 |
| Change in BAG (pretrained) | Future Naming (NEPSY-II) | -0.0280 (-0.17, 0.12) | 0.7020 | 1.0000 | -0.0036 | 0.0156 |
| Change in BAG (pretrained) | Future Inhibition (NEPSY-II) | 0.0376 (-0.11, 0.18) | 0.6073 | 1.0000 | -0.0031 | 0.0166 |
| Change in BAG (pretrained) | Future Switching (NEPSY-II) | 0.0021 (-0.14, 0.15) | 0.9772 | 1.0000 | -0.0043 | 0.0106 |

Table S14: Pretrained change vs. future results from GUSTO. p-values are bolded if less than $\alpha = 0.05$. **Key:** BAG – brain age gap; $\beta$ – standardized regression coefficient; $CI_L$ - lower limit of 95% confidence interval; $CI_U$ - upper limit of 95% confidence interval; $p$ – uncorrected p-value; $p_{corr}$ – corrected p-value; $\Delta R^2_{adj}$ - change in adjusted $R^2$ when adding variable of interest; $R^2$ - model coefficient of determination

| **Variable of Interest** | **Outcome** | $\beta$ $(\mathbf{CI_L}, \mathbf{CI_U})$ | **p** | $\mathbf{p_{corr}}$ | $\mathbf{\Delta R^2_{adj}}$ | $\mathbf{R^2}$ |
|---|---|---|---|---|---|---|
| Change in BAG (finetuned) | Future WCST Standard Score | -0.0091 (-0.16, 0.14) | 0.9052 | 1.0000 | -0.0045 | 0.0239 |
| Change in BAG (finetuned) | Future Naming (NEPSY-II) | 0.0158 (-0.14, 0.17) | 0.8413 | 1.0000 | -0.0041 | 0.0067 |
| Change in BAG (finetuned) | Future Inhibition (NEPSY-II) | 0.2006 (0.05, 0.35) | **0.0103** | **0.0411** | 0.0237 | 0.0400 |
| Change in BAG (finetuned) | Future Switching (NEPSY-II) | 0.1795 (0.03, 0.33) | **0.0221** | 0.0663 | 0.0181 | 0.0311 |

Table S15: Finetuned change vs. future results from GUSTO. p-values are bolded if less than $\alpha = 0.05$. **Key:** BAG – brain age gap; $\beta$ – standardized regression coefficient; $CI_L$ - lower limit of 95% confidence interval; $CI_U$ - upper limit of 95% confidence interval; $p$ – uncorrected p-value; $p_{corr}$ – corrected p-value; $\Delta R^2_{adj}$ - change in adjusted $R^2$ when adding variable of interest; $R^2$ - model coefficient of determination

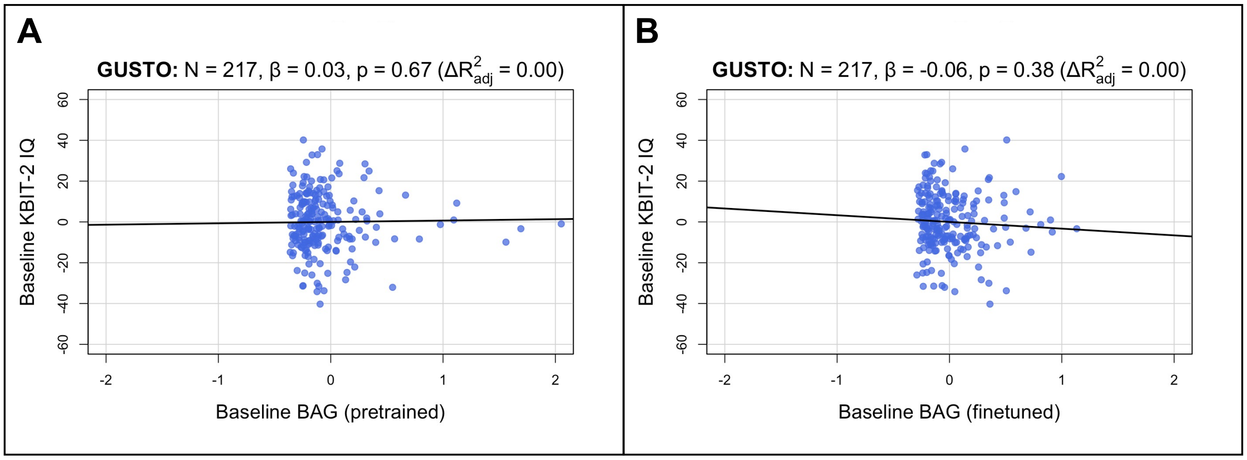


Figure S3: Baseline brain age gap is not associated with baseline IQ in children. Both brain age and IQ were measured cross-sectionally at 4.5 years old. **(A)** Partial regression plot with baseline BAG from the **pretrained** model. Estimated BAGs > 2 are not shown for visual clarity. No significant relationship is observed. **(B)** Partial regression plot with baseline BAG from the **finetuned** model. No significant relationship is observed. **Key:** N – number of participants; $\beta$ - standardized regression coefficient; p – p-value for variable of interest (x-axis); Multiple $R^2$ – coefficient of determination; GUSTO - Growing Up in Singapore Towards healthy Outcomes; BAG – Brain Age Gap; KBIT-2 - Kaufman Brief Intelligence Test Second Edition

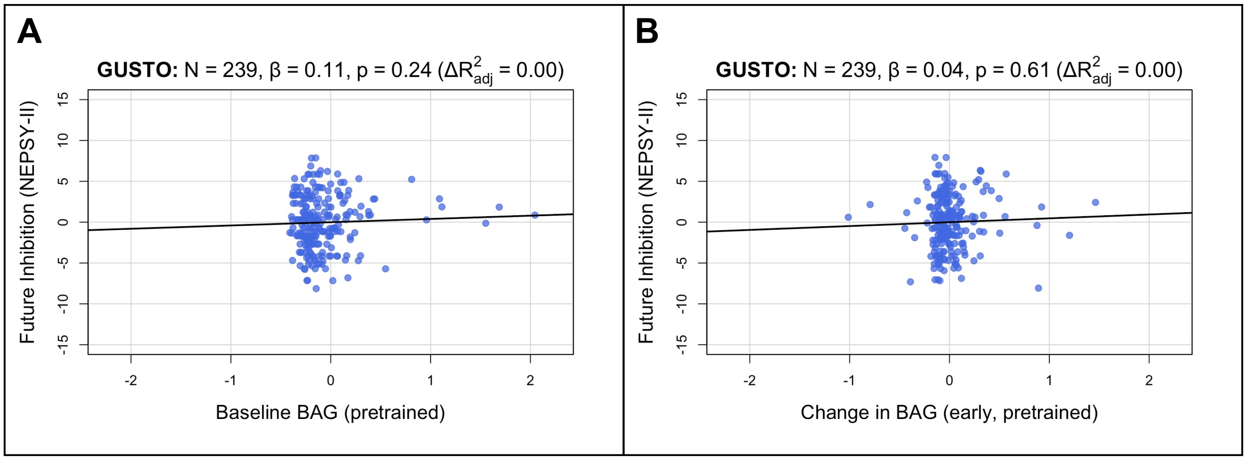


Figure S4: Brain age gap from the **pretrained** model is not associated with inhibition in **children**. Estimated BAGs > 2 are not shown for visual clarity. Compare to Figure 4 of the main text.

---

Figure S5 *(following page)*: **Pretrained** models focus on similar features as finetuned models in EDIS and SLABS, but not in GUSTO. Compare to Figure 5 of the main text.

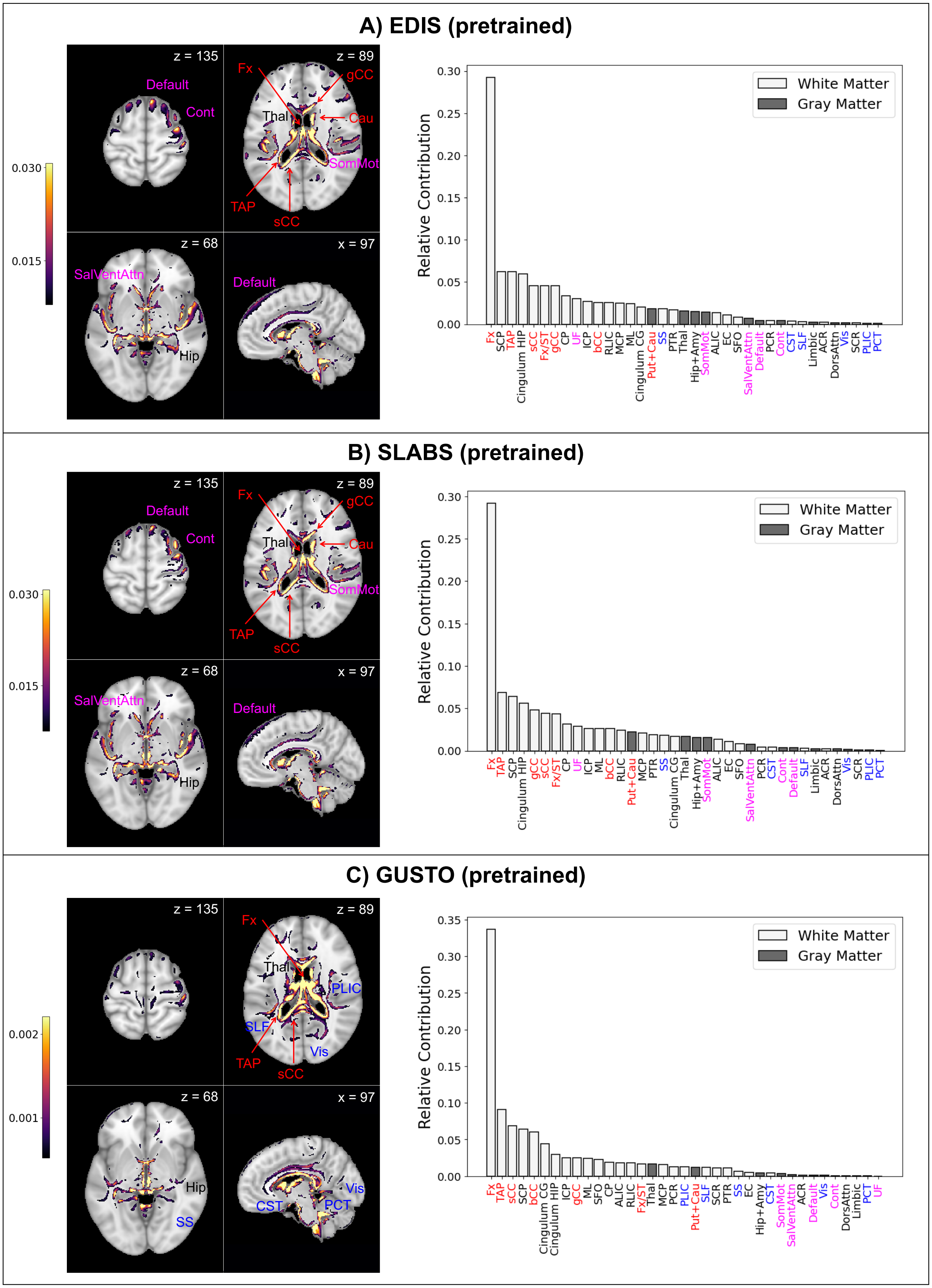

A) EDIS (pretrained)
z = 135
z = 89
Default
Cont
Fx
gCC
Thal
Cau
SomMot
TAP
sCC
z = 68
x = 97
SalVentAttn
Default
Hip
0.030
0.015
Relative Contribution
White Matter
Gray Matter
0.00
0.05
0.10
0.15
0.20
0.25
0.30
Fx SCP TAP Cingulum HIP sCC Fx/ST gCC CP UF ICP bCC RLIC MCP ML Cingulum CG Put+Cau SS PTR Thal Hip+Amy SomMot ALIC EC SFO SalVentAttn Default PCR Cont CST SLF Limbic ACR DorsAttn Vis SCR PLIC PCT
B) SLABS (pretrained)
z = 135
z = 89
Default
Cont
Fx
gCC
Thal
Cau
SomMot
TAP
sCC
z = 68
x = 97
SalVentAttn
Default
Hip
0.030
0.015
Relative Contribution
White Matter
Gray Matter
0.00
0.05
0.10
0.15
0.20
0.25
0.30
Fx TAP SCP Cingulum HIP gCC sCC Fx/ST CP UF ICP ML bCC RLIC Put+Cau MCP PTR SS Cingulum CG Thal Hip+Amy SomMot ALIC EC SFO SalVentAttn PCR CST Cont Default SLF Limbic ACR DorsAttn Vis SCR PLIC PCT
C) GUSTO (pretrained)
z = 135
z = 89
Fx
Thal
PLIC
SLF
TAP
sCC
Vis
z = 68
x = 97
Hip
SS
CST
PCT
Vis
0.002
0.001
Relative Contribution
White Matter
Gray Matter
0.00
0.05
0.10
0.15
0.20
0.25
0.30
0.35
Fx TAP sCC SCP bCC Cingulum CG Cingulum HIP ICP gCC ML SFO CP ALIC RLIC Fx/ST Thal MCP PCR PLIC Put+Cau SLF SCR PTR SS EC Hip+Amy CST SomMot SalVentAttn ACR Default Vis Cont DorsAttn Limbic PCT UF

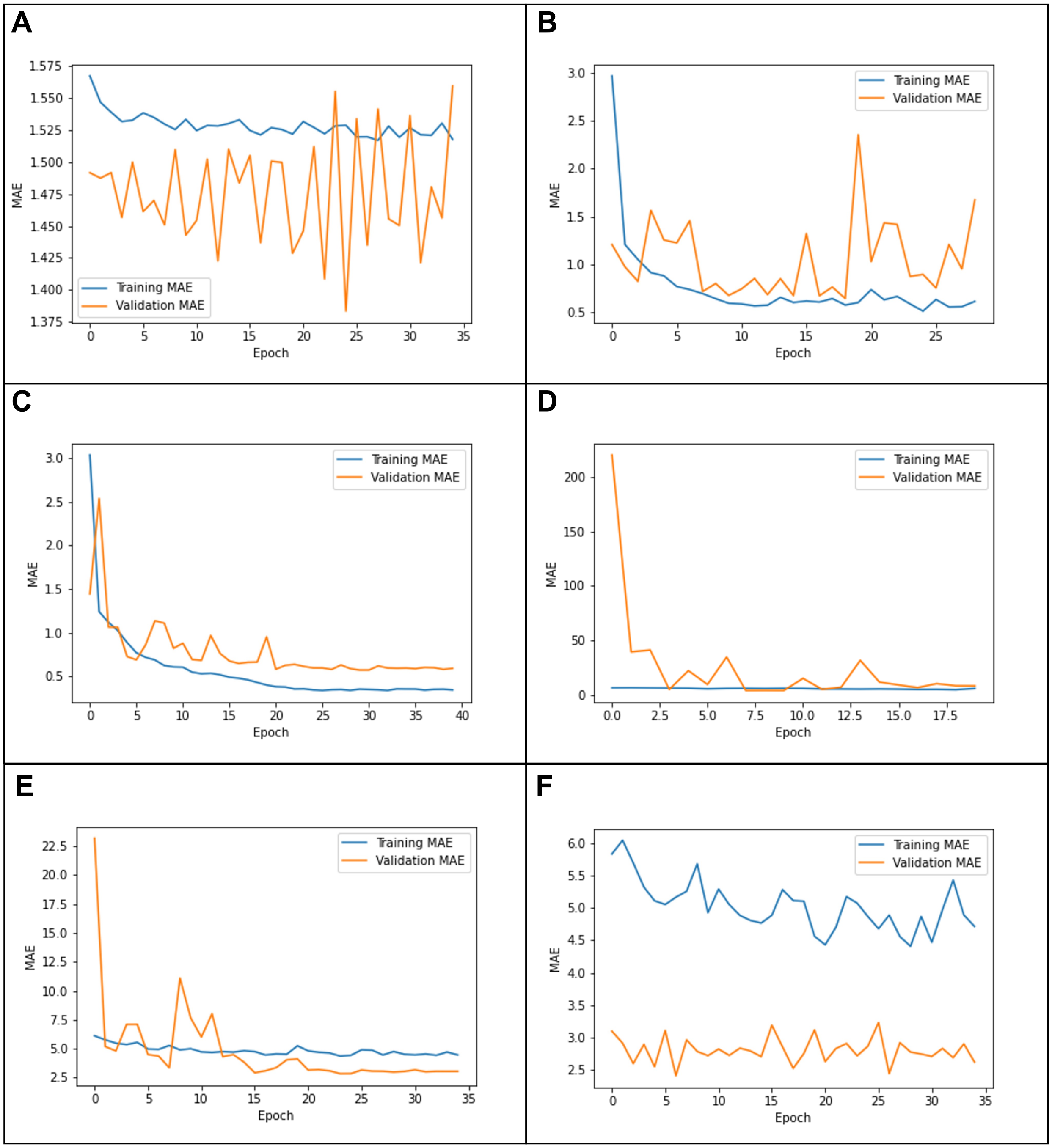

Figure S6: Example learning curves from **(A)** tuning the last layer only on GUSTO, showing underfitting; **(B)** tuning all layers on GUSTO, showing instability; **(C)** using a cosine learning rate decay, showing a good fit; **(D)** using the same parameters on EDIS, showing "forgetting;" **(E)** using a lower initial learning rate (1e-4) on EDIS, showing a better fit; and **(F)** using an initial learning rate of 1e-6, showing underfitting.
